# Quantitative Analysis of Miniature Synaptic Calcium Transients Using Positive Unlabeled Deep Learning

**DOI:** 10.1101/2024.07.04.602047

**Authors:** Frédéric Beaupré, Anthony Bilodeau, Theresa Wiesner, Gabriel Leclerc, Mado Lemieux, Gabriel Nadeau, Katrine Castonguay, Bolin Fan, Simon Labrecque, Renée Hložek, Paul De Koninck, Christian Gagné, Flavie Lavoie-Cardinal

**Affiliations:** CERVO Brain Research Center, Québec, Canada; Institute Intelligence and Data, Université Laval, Québec, Canada; Dunlap Institute for Astronomy and Astrophysics, University of Toronto, Toronto, Canada; David A. Dunlap Department for Astronomy and Astrophysics, University of Toronto, Toronto, Canada; Department of Biochemistry, Microbiology and Bioinformatics, Université Laval, Québec, Canada; Department of Electrical Engineering and Computer Engineering, Université Laval, Québec, Canada; Canada CIFAR AI Chair, affiliated to Mila; Department of Psychiatry and Neuroscience, Université Laval, Québec, Canada

## Abstract

Ca^2+^ imaging methods are widely used for studying cellular activity in the brain, allowing detailed analysis of dynamic processes across various scales. Enhanced by high-contrast optical microscopy and fluorescent Ca^2+^ sensors, this technique can be used to reveal localized Ca^2+^ fluctuations within neurons, including in sub-cellular compartments, such as the dendritic shaft or spines. Despite advances in Ca^2+^ sensors, the analysis of miniature Synaptic Calcium Transients (mSCTs), characterized by variability in morphology and low signal-to-noise ratios, remains challenging. Traditional threshold-based methods struggle with the detection and segmentation of these small, dynamic events. Deep learning (DL) approaches offer promising solutions but are limited by the need for large annotated datasets. Positive Unlabeled (PU) learning addresses this limitation by leveraging unlabeled instances to increase dataset size and enhance performance. This approach is particularly useful in the case of mSCTs that are scarce and small, associated with a very small proportion of the foreground pixels. PU learning significantly increases the effective size of the training dataset, improving model performance. Here, we present a PU learning-based strategy for detecting and segmenting mSCTs. We evaluate the performance of two 3D deep learning models, StarDist-3D and 3D U-Net, which are well established for the segmentation of small volumetric structures in microscopy datasets. By integrating PU learning, we enhance the 3D U-Net’s performance, demonstrating significant gains over traditional methods. This work pioneers the application of PU learning in Ca^2+^ imaging analysis, offering a robust framework for mSCT detection and segmentation. We also demonstrate how this quantitative analysis pipeline can be used for subsequent mSCTs feature analysis. We characterize morphological and kinetic changes of mSCTs associated with the application of chemical long-term potentiation (cLTP) stimulation in cultured rat hippocampal neurons. Our data-driven approach shows that a cLTP-inducing stimulus leads to the emergence of new active dendritic regions and differently affects mSCTs subtypes.

## 1 Introduction

Ca^2+^ imaging is widely used for the study of cellular activity modulations in the brain, enabling detailed monitoring and analysis of dynamic processes at various scales: from large neuronal networks down to individual synapses [1, 2]. Exploiting high-contrast optical microscopy approaches and fluorescent Ca^2+^ sensors, it offers insights into localized fluctuations of Ca^2+^ concentration within neurons [3, 4]. In the last decade, the development of a variety of sensitive Ca^2+^-sensors with increased brightness and improved detection kinetics has opened up new possibilities for the monitoring of dynamic processes occurring in small cellular compartments such as synapses [5, 6, 7]. For instance, Ca^2+^ imaging of miniature Synaptic Calcium Transients (mSCTs) has been studied to monitor the NMDA-receptor driven Ca^2+^ influxes generated by the spontaneous release of glutamate-containing presynaptic vesicles [3, 8, 9, 10, 11]. mSCTs are characterized by wide variability in their morphology and spatio-temporal propagation, but also by a generally small difference between peak and background intensity fluctuations [3]. This makes the task of selecting a robust and general method for automated mSCT analysis very challenging. Threshold-based methods relying on the extraction of regions of interest (ROIs) have been proposed for mSCT detection [12, 13, 14]. However, the accuracy of these methods is reduced for events that propagate or change size and shape over time. Thus, there is a need for automated analytical approaches to detect and segment small fluorescence signal fluctuations occurring at variable rates and locations, such as mSCTs [15]. Inclusion of noise-based thresholding was also proposed to improve the segmentation of small Ca^2+^ events [15]. Template-matching [16, 17, 18], spike deconvolution [19, 20], sequential Markov-Chain Monte Carlo (MCMC) [21] and approximate Bayesian inference [22] approaches have been developed to describe the underlying process by which spiking activity leads to fluorescence measurements. These methods focus on the kinetics of the detected Ca^2+^ transients, but fail to quantify Ca^2+^ concentration changes within small sub-cellular volumes, such as synapses, due to the inherently small changes in fluorescence intensity associated with these events. While user-defined analysis pipelines are still common, the bio-imaging field has seen a shift of paradigm in the tools developed for quantitative analysis. Indeed, further detection and segmentation performance gains can be obtained with the use of deep learning (DL) [23, 24, 25]. DL offers data-driven approaches to automatically learn features and patterns underlying the nature of structures and events present in a dataset. Consequently, these networks do not require to make *a priori* assumptions on either the regions of interest, model-based definitions of spatio-temporal features, or on parameter choices specific to a given task. They have been applied successfully to Ca^2+^ imaging detection tasks at the cellular level [26, 27, 28]. While those methods can be applied to high throughput ROI-based cell body detection of Ca^2+^ transients, they are not designed to detect and track sub-cellular events occurring at random locations and propagating over time such as mSCTs [15].

One limitation for the broad deployment of supervised DL-based Ca^2+^ imaging analysis is the need for large annotated datasets [29] to train the DL models. In many experimental contexts, including Ca^2+^ imaging, negative instances, corresponding to pixels that do not contain a Ca^2+^ event at a given time point, account for a large proportion of the image [30, 31]. This represents an additional challenge for the development of DL analysis strategies. In cases where Ca^2+^ transients (positive instances) are small and scarce, a large fraction of the foreground remains without annotation (unlabeled). Positive Unlabeled (PU) learning was proposed to leverage unlabeled instances for the training of deep neural networks [32, 33]. PU learning was first introduced in situations where the manual annotation of all negative instances is impractical, time-consuming, or too subjective to the given annotator [34]. During training, the PU DL model has access to positive instances and a predefined number of unlabeled instances. This significantly increases the size of the training dataset by providing additional information to the DL models and leveraging the abundance of unlabeled data. PU learning has already been applied with success to several bio-image analysis tasks [35, 36, 37].

Here, we present a PU learning-based strategy for the detection of mSCTs in cultured rat hippocampal neurons (Figure 1). The approach was developed to be applicable for the detection and segmentation of mSCTs characterized by a large diversity of signal amplitudes, frequencies, and spatial distributions. Our algorithm provides both mSCT detection and segmentation through the use of spatio-temporal DL models (3D U-Net [38] and StarDist-3D [39]). We introduce PU learning in the context of mSCTs detection, using foreground intensity as unlabeled instances and mSCTs as positive instances. In the absence of unlabeled instances for training, our algorithm is similar to that of **(author?)** [40], which implements a 3D U-Net for segmentation of low SNR Ca^2+^ events in full-frame confocal imaging data. We build upon this work and show that the addition of *unlabeled instances* to the training procedure leads to significant gains in the performance of the 3D U-Net. To the best of our knowledge, this is the first use of PU learning for Ca^2+^ imaging analysis. We leverage the trained DL model to create a multidimensional analysis framework, which allows the analysis of mSCT features associated with the application of a chemical long term potentiation (cLTP) inducing stimulus.

**Figure 1:**
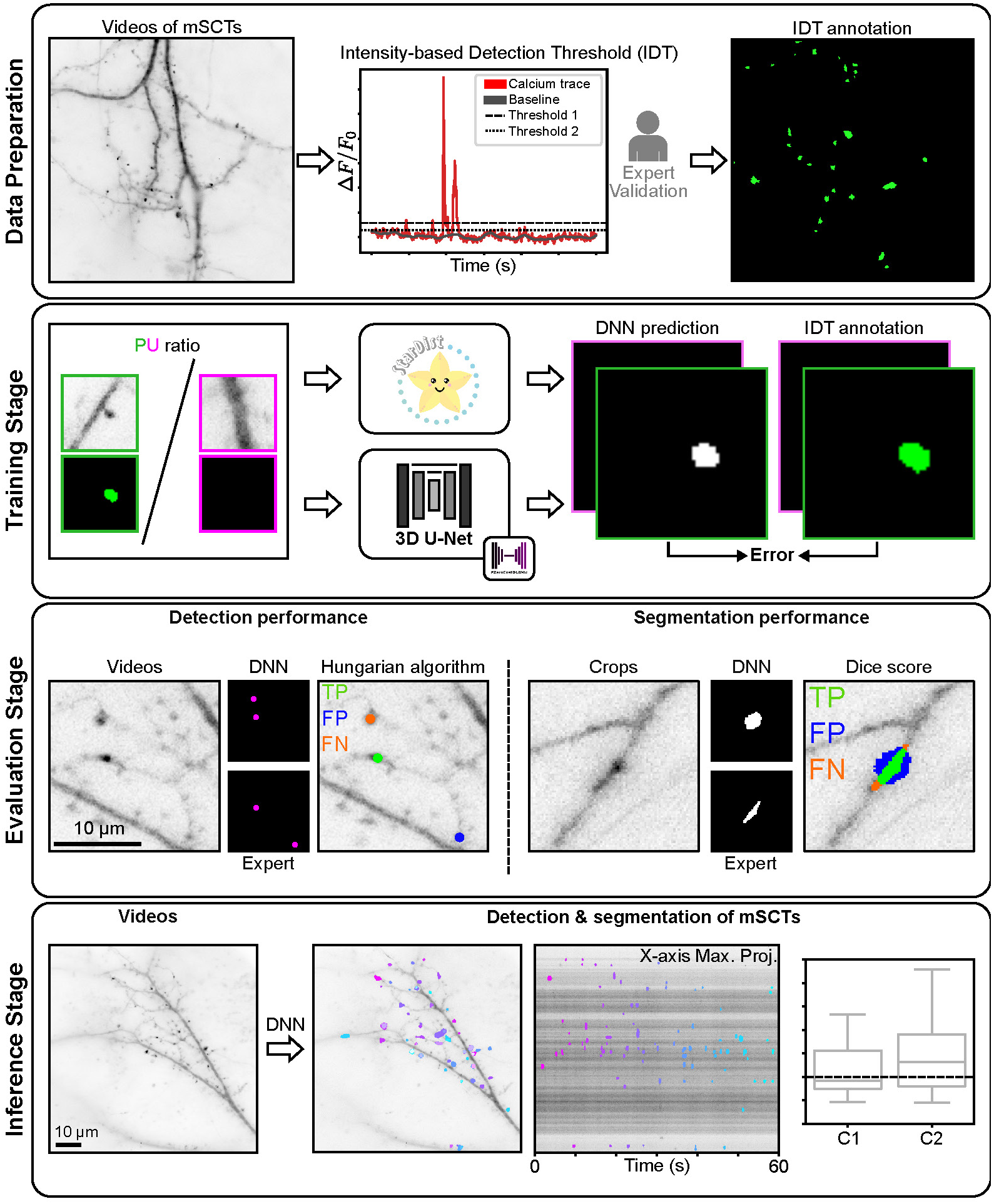
Deep learning algorithm workflow. **Data preparation:** Ca^2+^ imaging of the fluorescent protein GCaMP6f in cultured hippocampal neurons was performed with video microscopy. The transient localized fluorescent fluctuations are segmented using an intensity-based detection threshold (IDT) approach. Detected transients are validated by an expert to differentiate between foreground fluorescence signal fluctuation and mSCTs. **Training stage:** A DL model is trained on the dataset generated with the expert-validated IDT dataset with a PU learning scheme for automated detection and segmentation of mSCTs. **Evaluation stage**: Crops of the model predictions and expert ground truths are compared. Detection and segmentation performance are assessed according to a weighted centroid detection error and the dice similarity score, respectively. **Inference stage**: The trained DL model is used to infer segmentation masks from input Ca^2+^ imaging videos containing mSCTs. Maximum projections of the input video and predicted segmented events are shown. Predicted segmentation masks of mSCTs can be used for multidimensional feature analysis.

**Figure 2:**
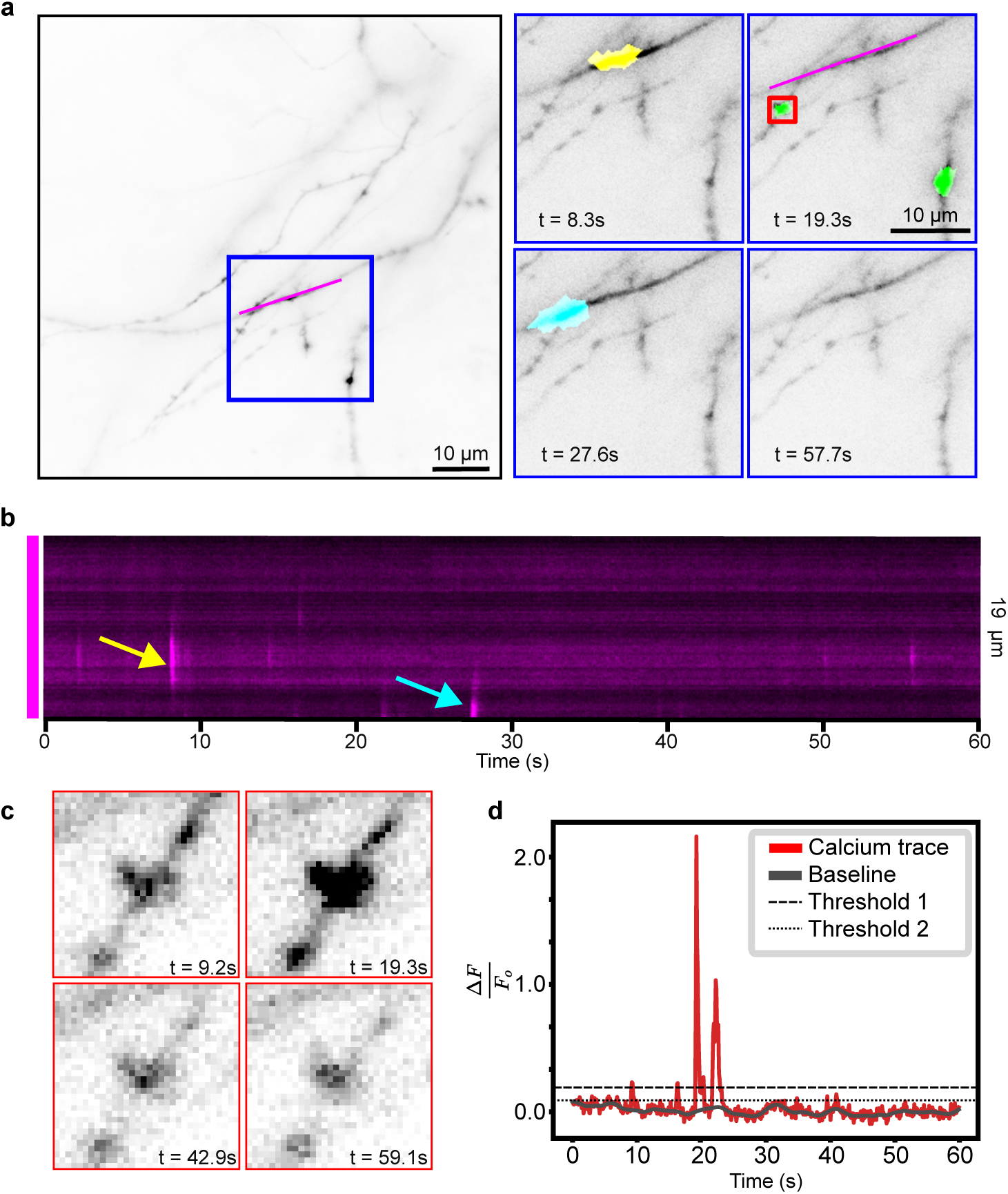
Video-microscopy of mSCTs. **a)** Left: Maximum projection of Ca^2+^ imaging video of GCaMP6f in cultured dissociated hippocampal neurons acquired on a custom-built TIRF microscope in oblique illumination mode. Right: Inset (blue square) highlighted in the left image of the full field of view at different time points. The yellow, green, and cyan regions in the insets are mSCTs detected by the IDT method. **b)** Kymograph of the dendritic region delimited by the magenta line in a). The arrows point to the events segmented in the top-left (yellow) and bottom-left (cyan) sub-region from a). **c)** (Left) Detected fluorescence signal measured at different time points for the highlighted region (red box) from the top right inset of panel a. **d)** Raw and baseline Ca^2+^ traces for the mSCT shown in c). IDT is based on two thresholds, for peak detection (Threshold 1) and for transient segmentation (Threshold 2)

## 2 Methods

### 2.1 Neuronal Cell Culture

Dissociated rat hippocampal cultures were prepared as described previously [41, 42]. In brief, hippocampi were dissected out of P1 rats, following approved protocols from the Université Laval animal care committee, and dissociated enzymatically (papain, 12U/ml; Worthington) and mechanically (trituration through Pasteur pipette). After dissociation, the cells were washed, centrifuged, and plated on poly-D-lysine–coated glass (18mm) coverslips at a density of *∼*67 500cells/mm^2^. Growth media consisted of Neurobasal and B27 (50:1), supplemented with penicillin/streptomycin (50 U/ml; 50 *µ*g/ml) and 0.5 mM L-glutamax (Invitrogen). Fetal bovine serum (2%; Hyclone) was added at time of plating. After 5 days, half of the media was changed without serum and with Ara-C (5 *µ*M; Sigma-Aldrich) to limit proliferation of non-neuronal cells. Twice a week thereon, half of the growth medium was replaced with serum- and Ara-C–free medium. For Ca^2+^ imaging, neurons were transfected with GCaMP6f-expressing plasmid (pGP-CMV-GCaMP6f) [6] using Lipofectamine 2000 (Invitrogen). For the plasticity experiments, neurons were transfected at day in vitro (DIV) 9-13 and imaged 24h later. For the generation of the training dataset for the detection and segmentation of mSCTs, videos from neurons imaged at DIV 7-17 were included.

### 2.2 Imaging solutions and pharmacology

The HEPES-buffered aCSF solution used for imaging contained (in mM) : 104 NaCl, 5 KCl, 10 Na-HEPES, and 10 Glucose. To initially maintain little or no NMDA receptor activity during the transfer of the coverslip to the microscope, this solution contained 0.6 CaCl_2_ and 5.0 MgCl_2_ (High-Mg^2^-aCSF). For mSCT measurements, Mg^2^ was removed, while 1.2mM Ca^2^ and 0.5*µ*M Tetrodotoxin (TTX, Alomone Labs) were added to the solution (0Mg^2^/TTX-aCSF). For chemical long term potentiation (cLTP) stimulation, the aCSF solution without Mg^2^ was supplemented with 0.2 mM glycine and 0.01 mM bicuculline (0 Mg^2^/Gly/Bic-aCSF) [43]. The osmolarity of every solution was maintained around 230 Osm, close to the neurobasal-based growth media value.

### 2.3 Live imaging and plasticity protocol

Neurons were imaged using a 100x 1.49NA immersion objective on an inverted Olympus IX-71, equipped with a TIRF illuminator, set on an oblique illumination mode to detect fluorescence signals across the z plane of dendrites with high contrast [44]. The microscope was equipped with a multi-valve perfusion system (VC3-8xG, ALA Scientifics instruments) and an inline heater (SF-28, Warner) so that neurons were continuously perfused with heated (32 °C) solutions. Transfected neurons were imaged with a Toptica iChrome MLE laser at 488 nm. GCaMP6f fluorescence emission was filtered through a Semrock, Brightline 520/35 cube placed before the objective and recorded on a Charge-coupled device (CCD) camera (Princeton ProEm 512B-FT) at a rate of 10 frames per second. The pixel size was 0.16 *µ*m. Image acquisition was performed using a custom written Matlab script on Matlab2007 [44].

For the mSCT plasticity experiments, neurons were transfected at DIV 9-13 and imaged 24h later. Coverslips were first transferred from the incubator to the perfusion chamber containing fresh High-Mg^2^-aCSF, to block the activity and prevent any cellular stress. The perfusion was switched for the imaging session to 0Mg^2^/TTX-aCSF and a baseline Ca^2+^-imaging video of 600 frames (60 seconds) was acquired. Immediately after, the perfusion was changed to 0Mg^2^/Gly/Bic-aCSF for 7 minutes for the cLTP stimulation. Neurons were not imaged during the cLTP stimulation to prevent photobleaching of the Ca^2^ sensor. Directly after the stimulation, the perfusion was switched back to 0Mg^2^/TTX-aCSF. A new video was acquired 10 min after the end of the stimulation. For the control condition, neurons were kept in the 0Mg^2^/TTX-aCSF throughout. For both conditions, the data shown were acquired 10 minutes apart.

### 2.4 Intensity-based Detection threshold (IDT) Annotator

We developed an intensity-based detection threshold (IDT) algorithm to detect and segment mSCTs while recording their temporal and spatial position. IDT is implemented in Matlab 2018 (Mathworks Inc., Natick, USA). It consists of 3 phases: i) data preprocessing, ii) automated detection and segmentation of mSCTs, and iii) manual verification of the detected Ca^2+^ events.

Acquired streams were initially aligned in x and y to the first frame of the video using the custom written Matlab code of **(author?)** [45]. Subsequently, the background noise was removed by manually choosing a region of interest in the background and by subtracting its average intensity from each frame. To detect the foreground, a median filter was applied and manual adjustment of an intensity-based threshold was performed. Ca^2+^ transient detection was performed pixel-wise from the foreground mask. For each pixel the fluorescence intensity was extracted and the baseline was corrected using asymmetric re-weighted penalized least square algorithm (arPLS) [46]. The fluorescence signal *F* was normalized to the baseline fluorescence *F*_0_ with [47]:

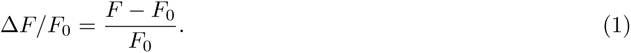

To identify real Ca^2+^ events from fluorescence foreground fluctuations, we first evaluated the noise of the baseline fluorescence trace. We compared the standard deviation (STD) of the signal while iteratively excluding the time points with the highest values, which correspond to Ca^2+^ events rather than noise fluctuations. When the STD between 2 consecutive iteration steps becomes similar, indicating that most of the Ca^2+^ events have been removed, we use the measured STD as an approximation of baseline noise fluctuations for subsequent steps. Ca^2+^ transients are detected with the Matlab function findpeaks. The first threshold is used to detect mSCTs and set at 4 *×* STD (detection threshold). A second threshold of 2 *×* STD is applied subsequently and used for the segmentation of the mSCTs (segmentation threshold). For each detected peak, in a window of 1500 ms before and after the maximal value, the segmentation threshold is used to determine the length of the mSCTs and generate a segmentation mask. The region properties of the objects are calculated using the regionprops function in Matlab. Detected events lasting only 1 frame or having a minor axis length *≤* 3 are removed. Pixels that have been identified as part of mSCTs are given the pixel value of 1. Following this first automated detection step, the user is asked to verify each detected event with the help of a graphical user interface (GUI). The GUI displays a small crop centered around each event (64 by 64 pixels, 12 frames) and the corresponding baseline-correction and detection on the normalized Ca^2+^ transients. In the GUI, the user inspects the detected events to determine which ones correspond to mSCTs and should be kept for future analysis. To facilitate the analysis, events detected at different time points but with weighted centroids closer than 2.55 µm are given the same region identifier (ID) and can be proofed at the same time. We noted that mSCTs which occur at a high frequency at the same position are often missed and that their segmentation typically lasts longer than 4.5 seconds. Hence, the user is asked to inspect these events with another GUI showing the mSCT (64 by 64 pixels, 12 frames) and the overlapped segmentation at the maximal peak intensity frame. The analysis time of a video strongly depends on the number of Ca^2+^ transients. On average we can find between 100 - 800 events and the analysis of such videos can require from 2 to 4 hours (automated and manual steps combined).

### 2.5 Dataset

The training dataset consisted of 14 Ca^2+^-imaging videos acquired on different neurons. To account for the very small size of the training dataset, we measured the averaged performance of each model trained with five independent random seeds and five distinct groups of 14 Ca^2+^-imaging videos annotated using the IDT algorithm. Positive crops (64 *×* 64 *×* 64 pixels) were generated for each transient detected by IDT and manually validated by an expert. For each subset, unlabeled crops were sampled from the foreground to obtain the desired PU ratio. Unlabeled crops from a smaller PU ratio are a subset of the next PU ratio, *e.g.* the unlabeled crops used to train the models with a PU ratio of 1-2 are contained in the ensemble of unlabeled crops used to train the models with a PU ratio of 1-4. The validation dataset was kept constant for all training and consisted of 15 independent Ca^2+^-imaging videos that were annotated with the IDT algorithm.

The testing detection dataset consisted of 10 independent Ca^2+^-imaging videos that were manually annotated by an expert at the peak intensity of the mSCT. The manual detection of mSCTs consisted of point annotations of the event centroids at the maximal Δ*F/F*_0_ frame.

The segmentation testing dataset consisted of manual segmentation masks obtained from two experts for 69 mSCTs of varied size, shape, and intensity. The masks were drawn on (32 *×* 32 pixels) pixel crops centred on the frame with maximal Δ*F/F*_0_ for each mSCT.

### 2.6 DL models

The DL models (StarDist-3D and 3D U-Net) were trained with the aligned and normalized version of the acquired Ca^2+^-imaging videos (baseline correction and Δ*F/F*_0_, see 2.4) and the corresponding IDT annotations. Training parameters varied for both models and are described below. All models were trained in PyTorch [48].

#### 2.6.1 3D U-Net

A modified 3D U-Net was implemented [38]. The encoder part of the architecture consisted of 5 layers of double convolution blocks followed by max pooling with a kernel size and stride of 2. The number of filters was doubled after each layer (8, 16, 32, 64, 128). Throughout the model, a kernel size of 3 with zero-padding and a stride of 1 were used. After each convolution, a batch normalization layer and leaky ReLU (0.02) were used. The bottleneck of the model consisted in two convolution layers, each followed by a ReLU activation. For the decoder part, features from lower layers were upscaled using transposed convolutions and concatenated with the information obtained from the skipping links located at the same depth. This was followed by a double set of convolution, batch normalization and ReLU. A final convolution layer projected the information to a single channel prediction and was followed by a sigmoid function.

The model was trained with the Adam optimizer, with default parameters and a learning rate of 0.0002, to minimize the mean quadratic error. The model was trained for 100k steps with a batch size of 128. Every 100 steps, the complete validation dataset was used to assess the generalization performance. Random crops and spatial flip were used during training for data augmentation. The model with the best performance on the validation test was kept for testing.

#### 2.6.2 StarDist-3D

We used the default implementation from [49, 39]. In brief, the StarDist approach proposes to localize objects of interest via star-convex polygons. More specifically, the StarDist method trains a simple U-Net architecture to densely predict object probabilities as well as star-convex polygons parameterized by the distances to object boundaries. Together, these two predictions produce an overcomplete set of candidate polygons for a given input image, after which the final result is obtained via non-maximum suppression (NMS) of the candidates.

The model was trained for 400 epochs with a batch size of 64 and 100 batches per epochs. The same data augmentation technique to the 3D U-Net was used during training.

### 2.7 Positive Unlabeled Learning

Typical supervised DL algorithms learn to represent data using raw inputs and their corresponding annotations [50]. A binary classification problem consists in training a model to discriminate two classes of events (e.g., positive and negative). Traditionally, supervised machine learning makes use of a dataset {(**x**_0_*, y*_0_), (**x**_1_*, y*_1_), …} where **x***_i_ ∈* R*^D^* is the input and *y_i_*, the associated annotation. In the case of the detection task we have *y_i_* ∈ {0, 1}, with the sample being positive when *y* = 1 and negative otherwise (*y* = 0).

In Positive Unlabeled (PU) learning [32], a binary variable *s* ∈ {0, 1} on whether an example was annotated or not is introduced in the training dataset {(**x**_0_*, y*_0_*, s*_0_), (**x**_1_*, y*_1_*, s*_1_), …}. When the annotation process does not make annotation errors, it implies that P(*y* = 1*|s* = 1) = 1. When *s* = 0, then the example can belong to both classes.

In some fields, such as biomedical imaging, deep learning algorithms can be difficult to implement due to the scarcity or cost of acquiring data belonging to the positive class, or label [51]. In this study, mSCTs account for a very small portion of the training dataset (*<* 0.05% of total pixels, or *<* 0.3% of foreground pixels). In a PU learning framework the network learns from positive (mSCTs detected with IDT) and *unlabeled* examples (extracted from the foreground). Negative and unlabeled examples differ in the assumption that each unlabeled example could belong to either the positive or negative class, albeit with a much larger chance of belonging to the negative class for very sparse and small events such as mSCTs. Consequently, PU learning is particularly appealing and efficient when unlabeled examples are available at low cost and in large quantities [32].

In our imaging dataset, mSCTs are the positive samples, and to construct a PU learning dataset, we sample regions along dendrites where no mSCTs were detected with IDT. As mSCTs account for such a small portion of the total video pixels, we can acquire U crops in large quantities. To better study the effect of the number of U crops on model performance, we first evaluate the performance of a model trained in a positive-only context (ratio 1-0). Both Stardist-3D and 3D U-Net are trained in a PU learning setting, with PU ratio ranging from 1-1 to 1-256.

### 2.8 Evaluation strategies

#### 2.8.1 Detection

To assess the detection performance of the DL models, we frame the evaluation as a linear assignment problem which we solve using the hungarian algorithm [52] and the *linear*_*sum*_*assignment* function from the Python library Scipy. The hungarian algorithm aims to assign elements from group A to elements from group B while minimizing the cost of assignment. In our case, the elements of groups A and B are the centroids of the model and ground truth segmentations, and the cost of assignment is defined as a distance metric (euclidean) between the centroids. We define a value of 6 pixels as the maximum distance for a possible assignment. The performance is evaluated on the detection testing dataset consisting of 10 videos in which the centroids of all mSCTs were identified by an expert. We identified true positives (TP), false positives (FP) and false negatives (FN) with 100 evenly spaced thresholds in the interval [0.0, 1.0], which allowed computation of recall and precision scores for IDT, StarDist-3D and 3D U-Net. We also compute the average precision (AP) of every DL model for a more concise report of performance. The AP metric is defined as:

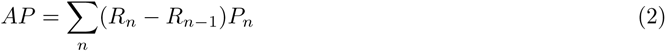

where *P_n_*and *R_n_*are the precision and recall at the *n^th^* threshold. Detection accuracy, recall and precision were also evaluated for different groups of event Δ*F/F*_0_. mSCT events were placed into 0.5 Δ*F/F*_0_ bin increments (except for the last bin which groups all events with Δ*F/F*_0_ *>* 3). The aforementioned metrics were computed for each bin, resulting in the matrices shown in Figure 3g. All detection results are averaged over the 25 effective training repetitions described in the *Dataset* section.

**Figure 3:**
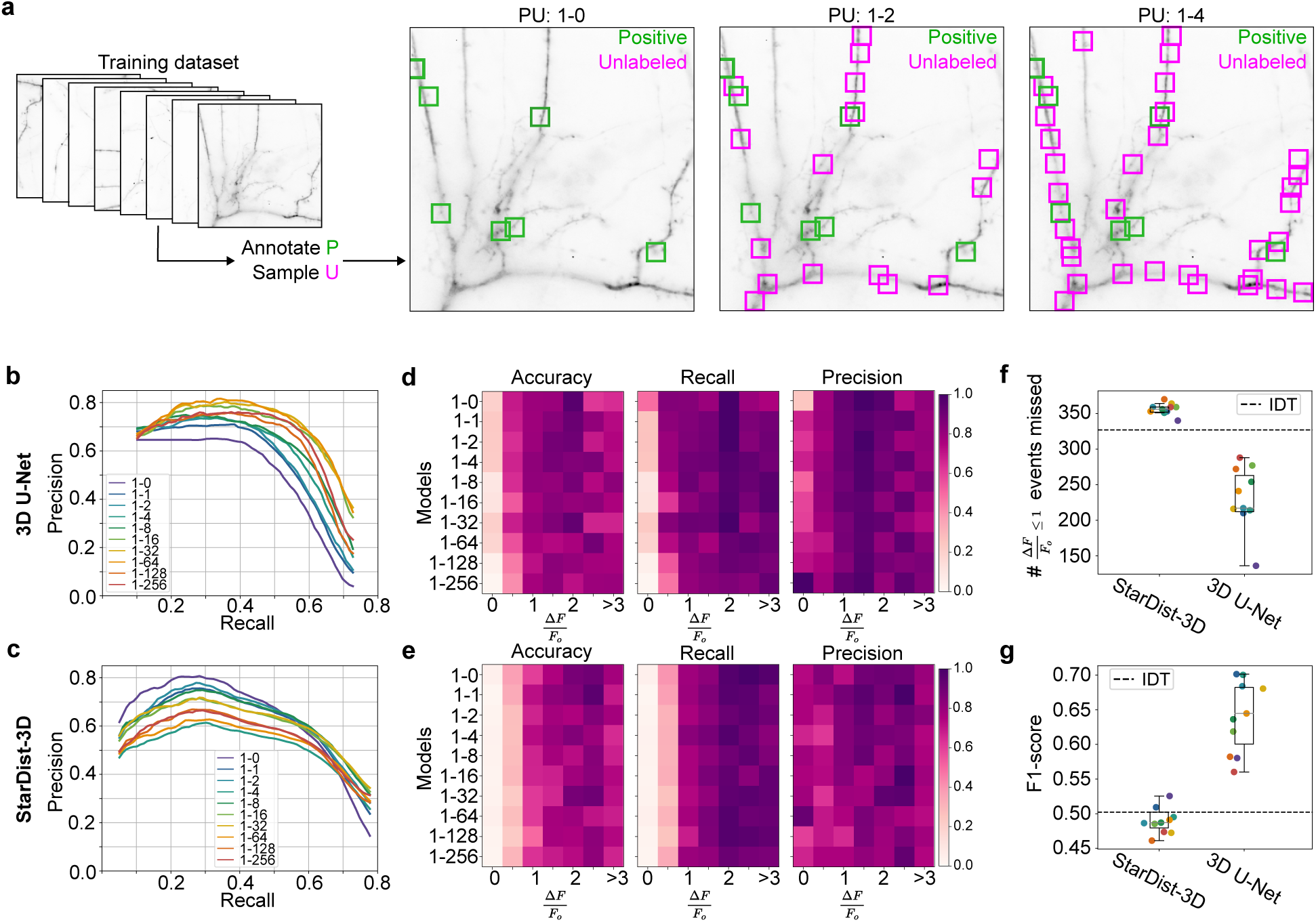
Detection of mSCTs with PU learning. **a)** The PU learning training dataset consists of 14 Ca^2+^-imaging videos in which mSCTs were labeled with IDT (positive instances). Unlabeled instances are sampled in each video of the training set according to the desired PU ratio. A maximum projection of a video is shown with an example of positive and unlabeled instances resulting from three different PU configurations. **b-c)** 3D U-Net (b) and StarDist-3D (c) precision-recall curves for the different PU ratio. The 1-0 configuration corresponds to the original dataset without unlabeled instances. **d,e)** Accuracy, recall, and precision of the different 3D U-Net models (d) and StarDist-3D (e) as a function of detected events’ Δ*F/F*0. **f)** Number of events with a Δ*F/F*0 *≤* 1 missed by each model. **g)** F1-score of the different models, using a detection threshold optimized on the validation set in the case of StarDist-3D and 3D U-Net. f,g) Color-coding of the data points corresponds to the different PU configurations for StarDist-3D and 3D U-Net shown in c.

#### 2.8.2 Segmentation

The segmentation performance of the DL models is reported as the dice similarity score [53] between the model’s binarized predictions and the ground truth segmentation obtained from the testing segmentation dataset. To measure intra-expert agreement, the same segmentation task was performed by experts on the testing dataset twice, with a two-week delay between annotation sessions. To measure inter-expert agreement, we compared the annotations from both experts in their first annotation session. Expert agreement was also measured using the dice similarity score. Considering that the annotations from expert 2 were in better agreement with IDT and in turn with the networks’ annotations, we used expert 2’s annotations as the segmentation performance of the models for different PU ratios (Figure 4c). We considered a model better than inter-expert level when it had a dice value at peak distribution density higher than that of the inter-expert distribution (gray vertical lines in Fig 4b). We opted for this definition rather than the distribution mean because of the double penalty that is being forced upon models which miss a large number of events (many false positives). Explicitly, models are being penalized twice on missed events; once in detection and once again in segmentation with a dice score of 0. This causes many models (especially for StarDist-3D) to show a bimodal dice score distribution with first mode near zero, which skews the distribution mean towards lower values. In contrast, experts were only shown frames displaying a mSCT, i.e., they could never be penalized for a missed event, and therefore do not exhibit this bimodal dice score distribution in their agreement.

**Figure 4:**
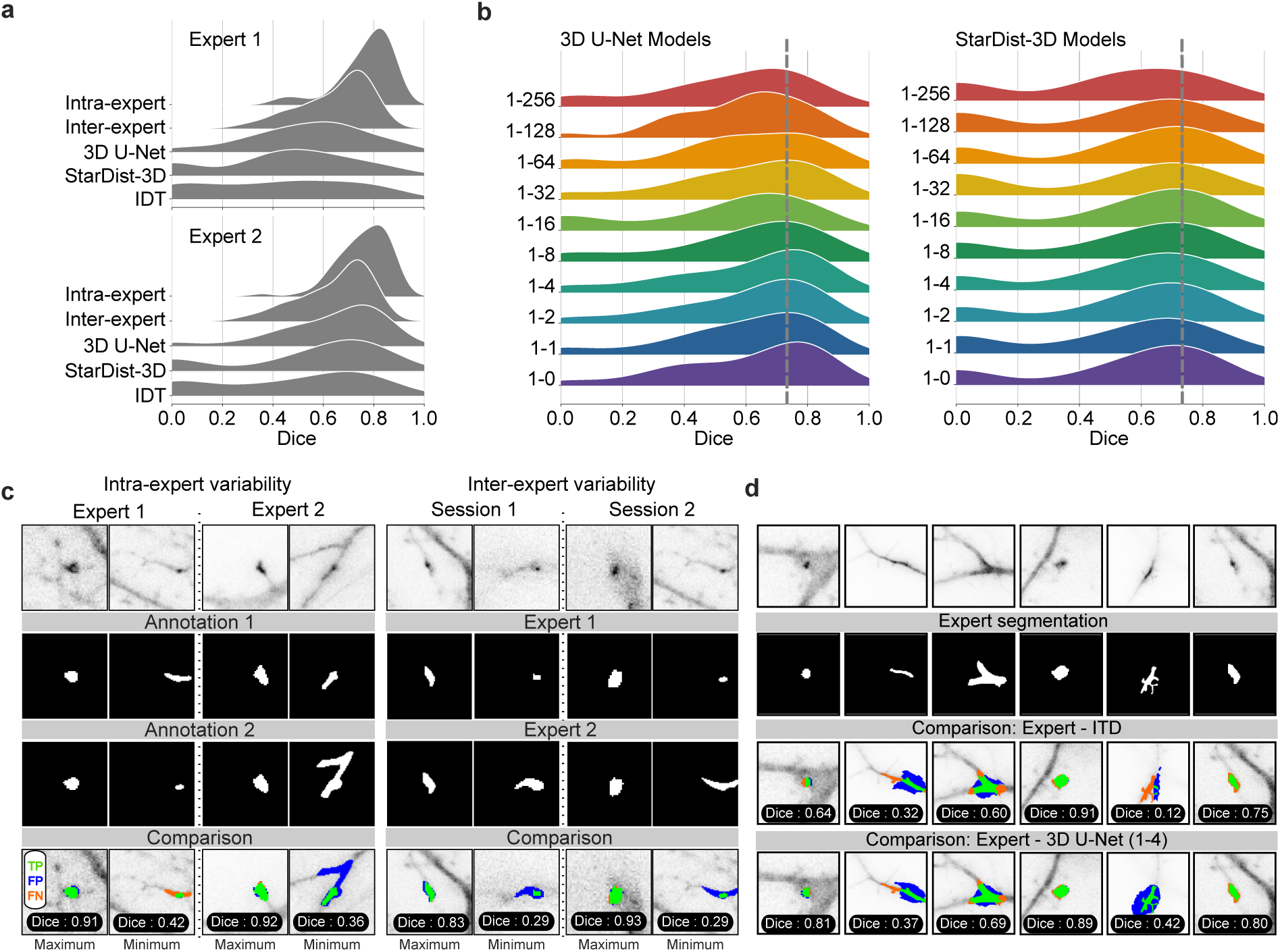
Segmentation of mSCTs. **a)** The segmentation performance of the models is measured on a testing dataset of 69 mSCTs (1 frame/mSCT) annotated by two experts. The segmentation masks of each mSCTs are compared to the one of each expert to account for inter-expert variability. Intra-expert agreement is evaluated using annotations from the same expert performed within a two week interval. The performance is reported for the DL models trained without negative instances (PU ratio 1-0). **b)** Segmentation performance of StarDist-3D and 3D U-Net for different PU configurations. Expert 2’s first annotation session is used as ground truth to measure the dice score. The 3D U-Net achieve an increased segmentation performance compared to the inter-expert agreement (gray dashed line) for PU ratios below 1-64. The best segmentation performance is obtained for a PU ratio of 1-4. **c)** Example annotations and resulting dice scores from the inter- and intra-expert studies. **d)** Example annotations (IDT) and predictions (3D U-Net with PU = 1-4) compared to the expert ground truths.

### 2.9 Multdimensional analysis

#### 2.9.1 Identification of mSCT subtypes

The mSCT subtypes are identified using the *K*-means algorithm. The number of subtypes (*K* = 5) is obtained from the Silhouette score [54] and the Elbow method [55], which find the optimal number of subtypes by computing at which value of *K* the Silhouette score is maximal and the Elbow method exhibits a linear decreasing trend with increasing number of subtypes (Supp. Fig. 3). More specifically, *K*-means is performed on the dataset *X ∈* R*^N×M^*, where *N* is the number of detected mSCTs and M is the number of features characterizing the mSCTs (note that these are manually defined, Supp. Tab. 2). Each detected mSCT is associated with its closest subtype using the pairwise euclidean distance between the averaged feature vectors of the subtypes and the feature vector of each mSCT.

#### 2.9.2 Determination of event categories

We determined event categories using the spatial coordinate of the segmentation masks. We can associate mSCTs occurring in the same region at different time points and classify them as: Silenced (S) when they are present only in the baseline video, New (N) for events detected only after the stimulus, and Recurrent (R) for events detected in the same region before (R_b_), and after (R_a_) the stimulus. A baseline event is considered recurrent when its weighted centroid is within 6 pixels (spatially) from an event detected in the second movie.

#### 2.9.3 Association of recurrent events

In Figure 5g, we associated the subtypes of recurrent baseline events R_b_ with the subtypes of their corresponding events detected after the cLTP stimulation R_a_. The colormap of the association matrix corresponds to the percentage of R_b_ events of a given subtype that were associated with a specific subtype R_a_.

**Figure 5:**
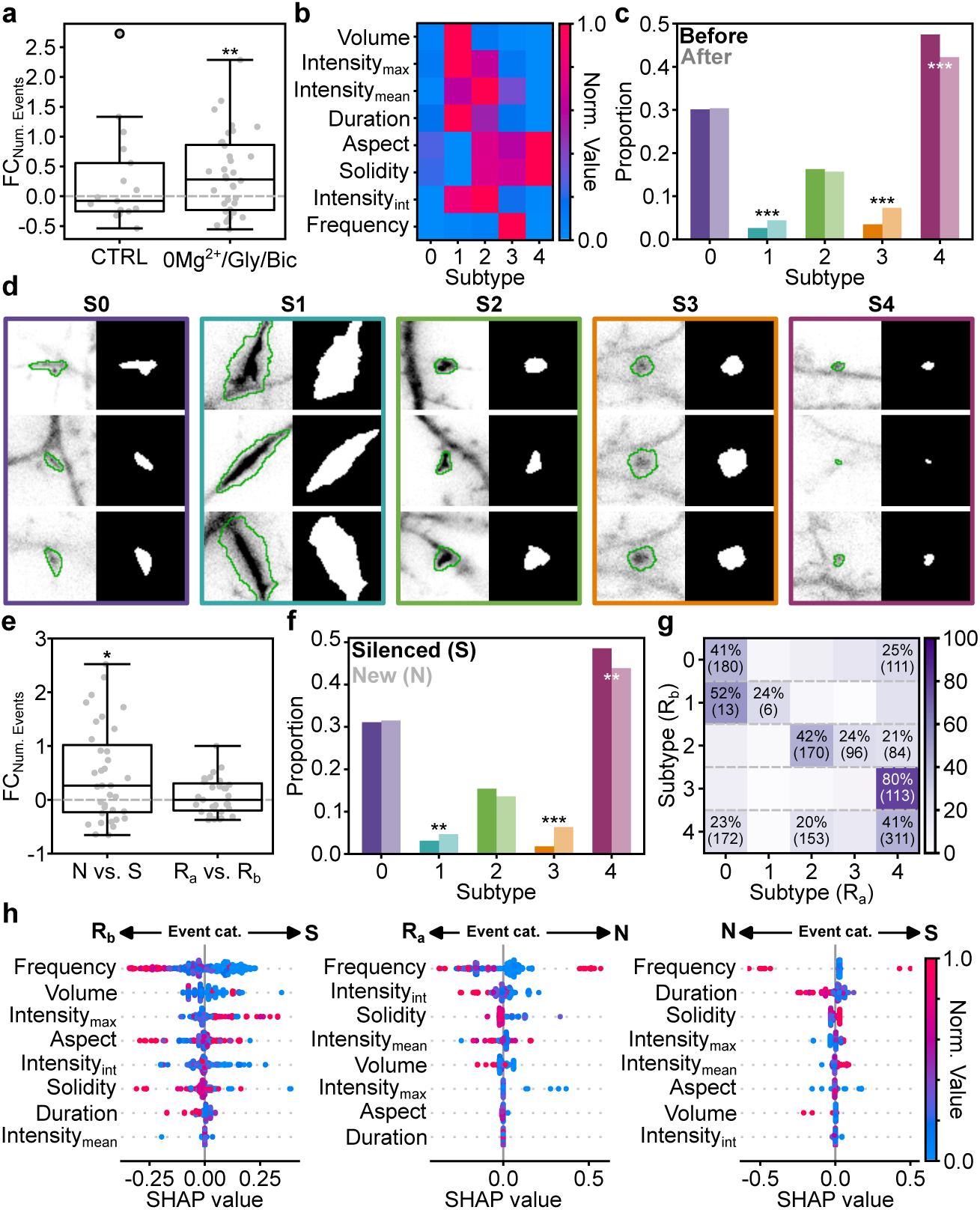
Quantitative analysis of mSCT features in a cLTP paradigm. **a)** The number of detected mSCTs is significantly increased following a cLTP-inducing stimulus (0Mg^2+^/Gly/Bic; Wilcoxon Signed Rank Test: *p*-value = 0.009), while it remains unchanged in the control condition (CTRL; Wilcoxon Signed Rank Test: *p*-value = 0.644). **b)** Averaged morphological and kinetic normalized feature values for each subtype obtained from *K*-means clustering (Supp Tab. 3 for normalization values). **c)** The proportion of events belonging to each subtype is modulated by a cLTP inducing stimulus. *χ*^2^ statistical analysis is performed (*p*-value = 1.63 *×* 10*^−^*^10^) followed by a post-hoc *χ*^2^ analysis test (S0: 8.24 *×* 10*^−^*^1^, S1: 7.70 *×* 10*^−^*^4^, S2: 5.35 *×* 10*^−^*^1^, S3: 3.83 *×* 10*^−^*^9^, S4: 2.13 *×* 10*^−^*^4^) **d)** Fluorescence signal and associated 3D U-Net segmentation for each mSCT subtype. (left) Image of detected transients at their maximal Δ*F/F*0 and the corresponding segmented borders (green overlay). (right) Segmentation mask predicted by the 3D U-Net for the frame with maximal Δ*F/F*0. Crops are 10.24 *µ*m *×* 10.24 *µ*m. **e)** The fold change in the number of new (N) events (FCNum. Events) is significantly increased compared to the number of silenced events following the cLTP stimulus (S, Wilcoxon Signed Rank Test: *p*-value = 0.02). The number of recurrent events is not significantly different before (R_b_) and after (R_a_) the stimulation (*t*-test: *p*-value = 0.30). **f)** The proportion of events belonging to each subtype is differently modulated for the silenced (dark) and new events (pale). *χ*^2^ statistical analysis is performed (*p*-value = 1.36 *×* 10*^−^*^10^) followed by a post-hoc *χ*^2^ analysis test (S0: 7.92 *×* 10*^−^*^1^, S1: 2.15 *×* 10*^−^*^2^, S2: 1.36 *×* 10*^−^*^1^, S3: 6.11 *×* 10*^−^*^11^, S4: 6.24 *×* 10*^−^*^3^). **g)** The subtypes of R_b_ events are associated with R_a_ events of different subtypes. Only *>* 20 % association are displayed. The raw number of events is shown in parenthesis. **h)** SHAP value obtained on the testing dataset from a Decision Tree Classifier trained to differentiate between event categories (R_a_ vs. S, R_b_ vs. N, and N vs. S) using the features extracted from the 3D U-Net segmentation mask for each detected mSCT (see Methods). Features are ranked according to their impact on the model output. Events are color-coded according to their normalized feature value (same normalization as in b).

#### 2.9.4 Feature-based Classification of mSCTs

We trained a decision tree on a binary classification task to assign mSCTs to distinct categories (Silenced, Recurrent, New) based on their features. The implementation from **(author?)** [56] was used. The hyper-parameters of the decision tree are given in Supp. Tab. 5. For each binary classification we used 70% of events for training and 30% for testing.

We computed the features with the most impact on the classifier’s predictions using SHapley Additive exPlanations (SHAP) values, which is a game theorety approach used to explain the output of machine learning model [57]. SHAP values give information regarding the contribution of each feature of the mSCT to the classifier prediction. The SHAP values predict how the machine learning model would perform with and without each specific feature to discriminate between the different mSCT categories, giving insights into the most important features to distinguish, for example, mSCTs detected only prior (silenced) or after (new) the cLTP simulation.

In Figure 5h, we plot the SHAP value for each event in the testing dataset, color-coded by their normalized feature values. The grey line corresponds to a SHAP value of 0, indicating no preference towards either class. Higher (or lower) SHAP values correspond to a greater influence on the model’s output. The SHAP values are arranged according to their distribution spread, with a larger spread indicating greater importance of this feature in the decision tree prediction. The color-coding of the events highlights the relationship between the value of each feature and its impact on the model’s prediction.

#### 2.9.5 Statistical analysis

The statistical difference for the fold change difference was tested using the Wilcoxon Signed Rank Test [58], which compares the distribution of differences between two observations and tests for the median of zero, rather than the *t*-test which uses the mean and would produce a different result if the distribution of distances was non-Gaussian, which we test for using the Shapiro-Wilk statistical test [59]. Otherwise, a *t*-test was used to determine the statistical significance. Points were considered as outliers and were discarded when they were 1.5 *× IQR* (interquartile range) away from the median.

The statistical difference between the proportion of subtypes was computed using the *χ*^2^ statistical analysis followed by a post-hoc *χ*^2^ analysis test.

## 3 Results

### 3.1 Intensity threshold-based detection of mSCTs

Ca^2+^ imaging was performed with video microscopy of cultured rat hippocampal neurons transfected with a genetically-encoded Ca^2+^ indicator (GCaMP6f) [6], using a TIRF microscope set in the oblique illumination mode [44]. To reveal mSCTs, the neurons were perfused with HEPES-buffered artificial cerebrospinal fluid (aCSF) lacking Mg^2+^ and supplemented with 0.5*µ*M Tetrodotoxin (TTX) to block action potentials. The localized transient changes in Ca^2+^ concentration, triggered by spontaneous release of glutamate and Ca^2+^ influx through NMDA receptors were monitored in proximal dendrites (Figure 2a-c). Videos of 60 s were acquired using a sampling frequency of 10 Hz. Lateral drift and background corrections were first applied to the raw videos (see Methods). The fluorescence signal *F* was normalized to the baseline *F*_0_ (Δ*F/F*_0_, see Methods) [47].

We developed an intensity-based detection threshold (IDT) approach for the detection and segmentation of mSCTs in the baseline-corrected dataset. We adopted a two-threshold approach to automatically detect and segment mSCTs in the Ca^2+^ imaging videos (Figure 2d), followed by manual inspection of the detected events. The first threshold (4 standard deviations (STD) above the baseline noise) was used to detect transients at their maximal Δ*F/F*_0_ value, while the second threshold (2 *×* STD) was used to generate a segmentation mask covering the entire duration of the mSCT (see Methods). A manual validation step of the detected transients was essential to confirm the detection results, as most mSCTs had a maximal Δ*F/F*_0_ *<* 1, making it difficult to distinguish real events from foreground fluorescence signal fluctuations using a fully-automated threshold-based approach. (Figure 1 – Data preparation). This manual step limited the number of false-positive detected events associated with noise and background fluctuations that were included in the final annotated IDT-dataset.

### 3.2 Detection of mSCTs with PU-learning

We used the manually curated IDT-annotated dataset of Ca^2+^ imaging videos as the ground truth to train the DL models in a supervised setting to automatically detect and segment mSCTs. The training sets consisted of 5 sets of 14 videos. The validation set included 15 annotated Ca^2+^ imaging videos. The detection performance was evaluated on a test set consisting of 10 Ca^2+^ imaging videos in which the *x, y, t* coordinates of the brightest point for all mSCTs were annotated by an expert. Manual annotation of the test set was necessary to overcome the lower detection accuracy of IDT for transients with a Δ*F/F*_0_ *≤* 1. To leverage the spatio-temporal dynamics of mSCTs, we chose 3D deep neural networks that are well established in the field of microscopy: 3D U-Net [38] and StarDist-3D [39]. The 3D U-Net was implemented in the `ZeroCostDL4Mic` standard [60] and is available for the community^1^. For StarDist-3D, we used the seminal implementation [49, 39]. Both of these DL models are trained to minimize the squared error (mean squared error loss) between their prediction and the corresponding IDT annotations (see Methods). To improve the DL models’ ability to differentiate between fluorescence signal fluctuations and mSCTs, we included in the training set unlabeled (U) foreground regions in addition to the positive (P) annotations obtained with IDT. We introduced a variable number of unlabeled instances to the training dataset (Figure 3a) to measure the impact of learning from unlabeled instances and validate the benefit of PU learning for a small data regime. The detection performance of the models trained with different PU ratios is compared using precision-recall (PR) curves (Figure 3b,c) to quantitatively evaluate the impact of leveraging unlabeled instances during training. For each PU ratio, the DL models were trained with 25 effective folds (5 seeds for 5 different positive-unlabeled training datasets - see Methods) and the averaged performance is reported. We apply the PU learning scheme by sampling unlabeled foreground instances to generate PU ratios ranging from 1-1 to 1-256. With the 3D U-Net, we observe maximal detection performance for a PU ratio of 1-64, after which the performances decrease, albeit remaining higher than the training scheme without unlabeled instances (ratio 1-0) (Figure 3b). For StarDist-3D, increasing the number of unlabeled instances negatively impacts its performance (Figure 3c). For PU ratios *<* 1 *−* 4, StarDist-3D shows a better performance than the 3D U-Net in terms of average precision (AP - see Methods). However, the 3D U-Net surpasses StarDist-3D for ratios above 1-4. Furthermore, the best 3D U-Net models (1-16, 1-32, 1-64, and 1-256) outperform the best StarDist-3D model in terms of AP, reaching a value of 0.59 for the 3D U-Net trained with a PU ratio of 1-64 (Supp. Tab. 4).

We then investigated the relationship between detection performance and mSCT fluorescence intensity (Figure 3d,e). Generally, mSCTs that are most difficult to distinguish from other fluorescence signal fluctuations are small and dim (Δ*F/F*_0_ *<* 1). These account for the majority of the events annotated by the expert in our testing dataset (Supp. Fig. 1). As expected, the performance (accuracy, recall and precision) of the 3D U-Net and StarDist-3D increases with increasing mSCT intensity, stabilizing for Δ*F/F*_0_ *>* 1 for all PU ratios (Figure 3d,e). The 3D U-Net shows an improved accuracy and recall over StarDist-3D for Δ*F/F*_0_ *<* 1 mSCTs (Fig. 3d,e). For the 3D U-Net, PU learning increases the precision at the cost of a small decrease in recall for PU ratios above 1-64. Training the 3D U-Net with unlabeled crops results in stricter models (better precision/fewer false positives; see Supp. Fig 2). This could be explained by the capability of the 3D U-Net to leverage information from unlabeled instances to refine its decision boundary between real mSCTs and foreground signal fluctuations. Figure 3f shows that the 3D U-Net misses fewer Δ*F/F*_0_ *<* 1 events than the IDT annotation method (black horizontal dashed line) and StarDist-3D. Similarly the 3D U-Net performs better than IDT and StarDist-3D in terms of F1-score for the detected mSCTs regardless of their Δ*F/F*_0_(Figure 3g, see Methods).

### 3.3 Segmentation of mSCTs

Accurate segmentation of mSCTs is important for the extraction of meaningful morphological (e.g. volume, area, shape) and kinetic (e.g. duration, frequency) features from the segmented fluorescence transients. We therefore evaluated the segmentation performance of our deep neural networks (across the same PU configurations as in Section 3.2) according to the dice similarity score [53]. A dice score of zero indicates no overlap and a dice score of unity indicates total overlap. To obtain segmentation ground truths, we asked two experts to manually annotate single frames of 69 mSCTs (see Methods). The type of events that were annotated by the experts varied in form, shape, and intensity. They included synaptic, dendritic, and out-of-focus events. Moreover, the annotation was performed by the same experts at two separate time points, to assess intra- and inter-expert agreement. Figure 4a shows the agreement results in terms of the dice score as well as the model performance when using either expert as ground truth. We first observe that expert 1 and expert 2 show similar levels of intra-expert agreement, with minimum mSCT segmentation dice scores of 0.42 and 0.36, respectively, means of 0.77 and 0.76, and maximums at 0.91 and 0.92. The inter-expert agreement values are lower, with minimum agreement between the two experts of 0.29, mean of 0.63, and maximum of 0.93. The rather low inter-expert mean dice score highlights the ambiguous nature of ground truths in bio-imaging datasets. These low agreement values suggest that the benchmark performance against which to compare the performance of the deep neural networks is better defined as the inter-expert level of agreement rather than a perfect dice score of 1. When comparing the segmentation performance of the DL models for different PU ratios (Figure 4b-d), we observe a slightly improved segmentation performance for the 3D U-Net compared to the inter-expert agreement for ratios below 1-64. Notably, ratios between 1-2 and 1-8 decrease the variability compared to the expert annotations: the number of segmentations with low dice scores below 0.5 decreases. As for the detection, PU learning does not improve the segmentation performance of StarDist-3D. Additionally, all StarDist-3D models show a bimodal dice distribution with first mode near zero. This indicates that StarDist-3D is unable to segment a significant number of ground truth events, which is evocative of the large number of missed events with Δ*F/F*_0_ *<* 1 (Figure 3f).

### 3.4 Multidimensional analysis of detected mSCTs

The presented detection and segmentation pipeline can serve as a valuable tool to extract morphology and kinetics features from mSCTs. We used the 3D U-Net trained with a PU learning ratio of 1-64 as it provides a good trade-off in detection and segmentation performance. The model was applied to the detection and segmentation of mSCTs on an unseen Ca^2+^ imaging dataset acquired on the same microscope. This dataset consists of paired videos of 600 frames, (see Methods) consisting of a baseline and a second video acquired 17 minutes after the end of the baseline video. In the control condition, the neurons were kept in a 0Mg^2+^/TTX solution throughout. We tested the effect of a cLTP stimulus on mSCTs using a bath application of a 0Mg^2+^/Gly/Bic solution during 7 minutes ([61, 62], see Methods) following the baseline video acquisition. The post-treatment video was acquired 10 minutes after the end of the stimulation, as in the control condition.

We observe a significant increase in the number of mSCT following the cLTP stimulus (Figure 5a). Seeing the heterogeneous distribution of morphology and kinetic features of mSCTs revealed by our quantitative analysis pipeline, we next extracted those features from the 3D U-Net’s mSCT segmentation masks (Supp. Table 2). From all extracted features, we performed a dependency analysis to identify the most relevant features and reduce interdependence between the variables. The segmented mSCTs were clustered using the *K*-means algorithm to identify mSCT subtypes in the dataset. The number of subtypes *K* = 5 is determined using the Silhouette score and the Elbow method (Supp. Fig. 3). The median value of the features for each subtype is shown in Figure 5b. To evaluate the effect of the cLTP treatment, we measured the proportion of mSCTs associated with each subtype before and after the cLTP stimulus (Figure 5c). We observe that the proportion of the most prevalent subtype (Subtype 4), which corresponds to round, short, and small Δ*F/F*_0_ Ca^2+^ events is significantly reduced after the stimulus. Meanwhile, the proportion of high frequency Ca^2+^ events (Subtype 3), as well as of large events associated with a high Δ*F/F*_0_ (Subtype 1), is significantly increased by the cLTP stimulus. Figure 5d shows examples of the different mSCT subtypes together with the 3D U-Net’s segmentation mask that were used to perform the feature analysis.

Using the spatial coordinate of the segmentation masks, we can associate mSCTs occurring in the same region at different time points and classify them as : Silenced (S) when they are present only in the baseline video, New (N) for events detected only after the stimulus, and Recurrent (R) for events detected in the same region before (R_b_), and after (R_a_) the stimulus. The increase of events following the cLTP stimulus (Figure 5a) is supported by the emergence of new active regions (Figure 5e). We next compared the proportion of subtypes between silenced and new events (Figure 5f). The measured proportions closely follow the overall proportion that was measured when combining all events (Figure 5c). To measure whether the treatment impacts the features of recurrent events differently, we associated the subtype of each R_b_ event with that of R_a_ events. As shown in Figure 5g, associated mSCTs do not necessarily share the same subtype before and after the stimulus but stronger correspondence between specific subtypes before and after the stimulus indicates that the modulation of recurrent events is dependant on the features of the Ca^2+^ event before the cLTP treatment.

We next evaluated the importance of each feature to discriminate between the different mSCT classes (S, N, R_a_, R_b_). We trained a decision tree classifier to differentiate mSCTs according to their classes, and computed the SHAP values to reveal the most important features leading to the predictions. SHAP values are also used to assess the impact that feature values have on the predicted outcome (in this case the classification task). In the SHAP value plot (Figure 5h), the features are presented in decreasing order of importance (most important = top-most feature), and the feature values (blue data point = low value, red data point = high value) are ordered along the x-axis according to whether they are likely to cause the model to predict that a mSCT belongs to one class or to another class. From Figure 5h, one can see that Silenced mSCTs and New mSCTs are principally differentiated from Recurrent mSCTs by their frequency. The second most important feature indicates that smaller events in the baseline videos are generally associated with Silenced (S) mSCTs compared to Recurrent (R_b_) mSCTs. A lower integrated intensity is beneficial to differentiate between New (N) and Recurrent (R_a_) mSCTs in the videos acquired after the cLTP stimulus. The SHAP values also indicate that the duration of the event is useful in discriminating between S and N events, but has only little impact for the other classes (Figure 5h). Our automated detection and segmentation approach with the 3D U-Net, combined with a multidimensional feature analysis of mSCTs showed that the increase in the event number measured following a cLTP-inducing stimulus is supported by the appearance of new active regions. Those new regions have distinct features, among which the proportion of events with high frequency is significantly increased, while small and short events are reduced following the stimulus.

## 4 Discussion

Through the application of PU learning to the quantitative analysis of Ca^2+^ imaging data, we have provided accurate detection and segmentation of mSCTs in fluorescence microscopy videos. We could extract signal dynamics and meaningful morphological features from these heterogeneous, low density (both spatial and temporal), low intensity, and propagating events. We have demonstrated that the use of negative instances in a PU learning framework can improve the performance for the detection of low intensity Ca^2+^ that are missed by other approaches, including other DL methods using solely positive examples for training. The improved performance observed with the 3D U-Net models was achieved simply by adding low-cost, quickly-sampled unlabeled instances to the existing positive instances. The training pipeline is implemented in an easy-to-use ZeroCostDL4Mic Jupyter notebook enabling straightforward training and fine-tuning of the models on new datasets, and specific use cases [60].

We further showed that our approach allows the extraction of features from the segmentation masks to measure the effect of a cLTP stimulus on mSCT morphologies and dynamics. The Ca^2+^ events extracted from the 3D U-Net detection and segmentation pipeline could be clustered into 5 main subtypes, that are differently modulated by the cLTP protocol. Having access to precise spatio-temporal localization of the mSCTs revealed that such cLTP stimulus leads largely to the emergence of new active regions, rather than an increase in the frequency of events within regions already active prior to the stimulus. This result may reflect a form of plasticity that is part of the mechanisms supporting long term potentiation of synaptic transmission [63, 64]. This potentiation may reflect changes in release probability in some axonal terminals [65, 66] (refs), increase recruitment of Ca^2+^ permeable ion channels in post-synaptic regions [67, 68], or an increase coupling between receptor activation and Ca^2+^ store release [69, 4] (refs). The ability to identify and analyze localized regions of activity will allow to dissect these various mechanisms and correlate them with molecular trafficking of proteins, such as the Ca2+/Calmodulin-Protein kinase II [70], or cytoskeletal remodelling of dendritic actin filaments [61].

Using the segmentation masks, we could also train a decision tree classifier to discriminate between events depending on the moment they were detected (before or after stimulation). SHAP values provided better insights into which features predominantly influence the model’s predictions, revealing that different features are used to distinguish recurrent events from new and silenced events.

The presented method should be readily applicable to many other small datasets of Ca^2+^ imaging, in which the events of interest are sparse and heterogeneous, providing insight into different applications and biological inquiries. We also showed that it can be applied to datasets with missing ground truth annotations, and still provide improvement on these annotations through the PU training of the networks, enabling better modeling of the boundaries between real Ca^2+^ transients and foreground signal fluctuations. While we have considered a simple formulation of PU learning, it could also be of interest to consider more sophisticated definitions such as in **(author?)** [33] on variety of bio-imaging datasets. Integration of the DL models into an accessible platform such as *ZeroCostDL4Mic* [60] should also facilitate the application of the proposed framework to new datasets or using other annotation methods as ground truths for training.

## Acknowledgments

We acknowledge the Neuronal Cell Culture Platform of the CERVO Brain Research Center for the preparation of the dissociated hippocampal cultures. We thank Annette Schwerdtfeger for careful proofreading of the manuscript. Funding was provided by grants from the Natural Sciences and Engineering Research Council of Canada (NSERC) (RGPIN-2019-06704 to F.L.C., RGPIN-2017-06171 to P.D.K. RGPIN-2018-05750 to R.H.), Fonds de Recherche Nature et Technologie (FRQNT) Team Grant (2021-PR-284335 to F.L.C, C.G. and P.D.K.), the Canadian Institute for Health Research (CIHR) (471107 to F.L.C. and P.D.K.), a New Frontiers in Research Fund Exploration Grant (NFRFE-2020-00933 to F.L.C., C.G. and R.H.), and the Neuronex Initiative (National Science Foundation 2014862, Fonds de recherche du Québec - Santé 295824 to F.L.C. and P.D.K.). R. H. acknowledges support from CIFAR, the Azrieli and Alfred. P. Sloan foundations and the Connaught Fund. C.G. is a CIFAR AI-Chair and F.L.C. is a Canada Research Chair Tier II. A.B. was supported by a scholarship from NSERC. F.B. was supported by scholarships from FRQNT, NeuroQuébec, and the Québec Bio-Imaging Network. A.B., F.B., and T.W. were awarded an excellence scholarship from the FRQNT strategic cluster UNIQUE.

## Author contributions

F.B., A.B., G.L., R.H., C.G., and F.L.C. designed the deep learning analysis strategy. T.W., M.L., and P.D.K. designed the Ca^2+^-imaging experiments. T.W. performed the Ca^2+^-imaging experiments. S.L. developed the Ca^2+^-imaging acquisition software. T.W., M.L., S.L. and G.N. developed the IDT approach. A.B., F.B., and F.L.C. designed the multidimensional analysis approach. T.W. and G.L. designed the user-study. T.W., G.L. and F.L.C. annotated the testing datasets. K.C. and B.L.F. tested implementations of IDT and PU learning strategies. F.B. and A.B. implemented the deep learning models, performed the PU learning experiments, and analyzed the results. F.B., A.B. and F.L.C wrote the manuscript. R.H., P.D.K., C.G. and F.L.C co-supervised the project.

## Competing interests

The authors declare no competing interests.

## Data availability

All datasets used to train the models and the corresponding acquired images are available for download at

https://s3.valeria.science/flclab-calcium/index.html.

## Code availability

All code used in this manuscript are open source and available at : https://github.com/FLClab/Calcium-Analysis.

## Footnotes

1 https://github.com/FLClab/Calcium-Analysis

## Notes

https://s3.valeria.science/flclab-calcium/index.html

### Competing Interest Statement

The authors have declared no competing interest.

