## Supplementary Figures and Tables for "Quantitative Analysis of Miniature Synaptic Calcium Transients Using Positive Unlabeled Deep Learning"

### Supplementary Material

#### Tables

| Model | TP % | FP % | FN % |
|---|---|---|---|
| IDT | 33.0 | 14.3 | 52.7 |
| StarDist 1-0 | 35.6 | 7.9 | 56.5 |
| U-Net 1-0 | 36.4 | 50.4 | 13.2 |
| U-Net 1-64 | 49.8 | 15.7 | 34.5 |

Supplementary Tab. 1: Proportion of true positives, false positives and false negatives per model.

| Feature | Description |
|---|---|
| Volume | Volume of an event |
| $\text{Intensity}_{\text{max}}$ | Maximum $\Delta F/F_0$ of the entire event |
| $\text{Intensity}_{\text{mean}}$ | Mean $\Delta F/F_0$ of the entire event |
| Duration | Length in time of the event |
| Aspect | Aspect ratio of the event calculated as $\frac{\text{minor}}{\text{major}}$ |
| Solidity | Solidity of an event |
| $\text{Intensity}_{\text{int}}$ | The integration of the averaged intensity profile in time of an event |
| Frequency | Number of events at the event $(x, y)$ location |

Supplementary Tab. 2: Shape and intensity features of events. Most features are extracted by using `regionprops` from Scikit-Image [71].

| Feature | minimum | maximum |
|---|---|---|
| Volume (Voxels) | 112 | 6777 |
| $\text{Intensity}_{\text{max}}$ | 1.04 | 5.55 |
| $\text{Intensity}_{\text{mean}}$ | 0.45 | 1.03 |
| Duration | 4 | 10 |
| Aspect | 0.16 | 0.49 |
| Solidity | 0.73 | 0.89 |
| $\text{Intensity}_{\text{int}}$ | 0.96 | 3.37 |
| Frequency | 0 | 13 |

Supplementary Tab. 3: Values used to normalize the features in Figure 5b,h

|  | 3D U-Net | StarDist-3D |
|---|---|---|
| PU ratio | Average Precision | |
| 1-0 | 0.39 ± 0.06 | 0.50 ± 0.03 |
| 1-1 | 0.44 ± 0.08 | 0.48 ± 0.03 |
| 1-2 | 0.46 ± 0.07 | 0.52 ± 0.04 |
| 1-4 | 0.49 ± 0.08 | 0.34 ± 0.04 |
| 1-8 | 0.51 ± 0.09 | 0.53 ± 0.04 |
| 1-16 | 0.56 ± 0.12 | 0.49 ± 0.04 |
| 1-32 | 0.58 ± 0.10 | 0.52 ± 0.04 |
| 1-64 | **0.59 ± 0.08** | 0.40 ± 0.04 |
| 1-128 | 0.51 ± 0.11 | 0.45 ± 0.03 |
| 1-256 | 0.54 ± 0.07 | 0.46 ± 0.03 |

Supplementary Tab. 4: Average precision of the different models averaged over 25 effective seeds with standard deviation shown. In bold is the top performing model (3D U-Net with PU ratio of 1-64).

| Hyper-parameter | Value |
|---|---|
| `min_sample_split` | 2 |
| `max_depth` | 5 |
| `class_weight` | balanced |

Supplementary Tab. 5: Hyper-parameters of the Decision Tree Classifier that were used to classify the type of events ($R_b$ vs. S, $R_a$ vs. N, and N vs. S).

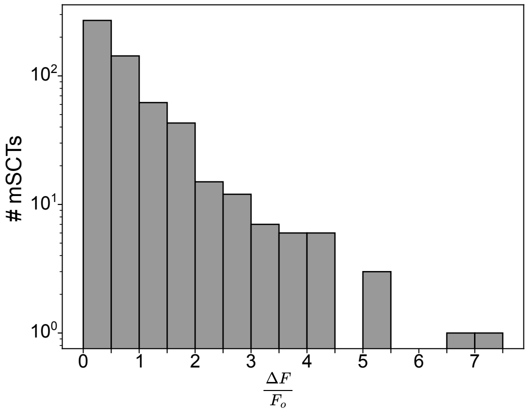


Supplementary Fig. 1: Histogram of the number of ground truth MSCTs as a function of their $\Delta F/F_0$.

### Figures

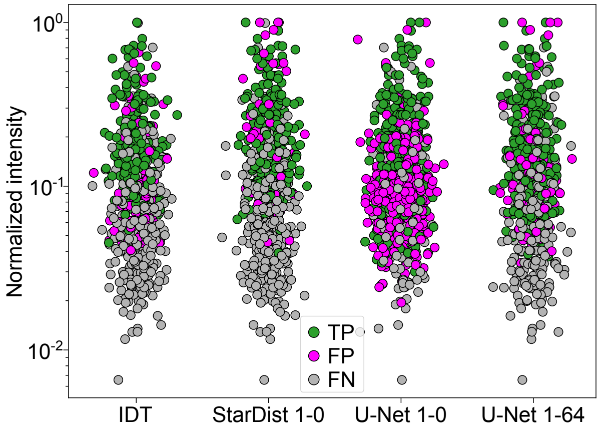


Supplementary Fig. 2: Events detected by the models plotted as a function of their intensity and their detection type (True Positive, False Positive or False Negative).

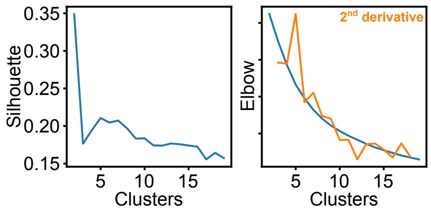


Supplementary Fig. 3: Silhouette and Elbow score as a function of the number of clusters from $k$-means for the segmentation of mSCTs in the cLTP condition. For the Elbow method, the $2^{nd}$ is presented to show the highest deflection point. Both methods agree for 5 clusters.
